# Shallow environmental gradients can cause range margins to form

**DOI:** 10.1101/2022.03.19.484973

**Authors:** Matteo Tomasini, Martin Eriksson, Kerstin Johannesson, Marina Rafajlović

## Abstract

One hypothesis invoked to explain limits to species’ ranges is a mismatch in environmental conditions between the central and marginal areas of species’ ranges. Low population size at the margins causes genetic drift to outplay selection locally, and limits the accumulation of genetic variance, so that adaptation is hindered locally. Earlier theoretical work shows that, for a population expanding over a spatially heterogeneous environment without any geographical barriers, adaptation will fail abruptly and sharp range margins will establish only when the underlying environmental conditions change more and more severely across space, whereas an environment changing constantly will result either in infinite expansion or rapid global extinction. Here, we extend this “steepening-gradient hypothesis” to encompass situations when multiple (up to three) environmental factors impose selection on separate adaptive traits. We show that multiple selection gradients steepen each other and that it is sufficient that just one of the gradients steepen in space for sharp range margins to form. This is true even if this gradient is shallow throughout the realised range. Thus, despite its detrimental role in forming range margins, it could be overlooked in field studies. Finally, by decomposing an environmental gradient to selection on two (or three) adaptive traits, we show that a population can withstand harsher environmental conditions than when selection acts on one adaptive trait alone. This finding argues for the evolution of novel traits in harsh environments.

## Introduction

Populations’ range margins can form due to discontinuities in the environment, such as geographical barriers (*e*.*g*. mountain ranges, rivers) [Holt, 2003]. However, sometimes range margins seem to occur abruptly along smooth environmental gradients, in the absence of any obvious geographical barriers [Bridle and Vines, 2007]. A question that has puzzled ecologists and evolutionary biologists for a long time, is whether range margins can arise solely as a result of a failure to adapt along spatially changing environmental conditions [Mayr, 1963, MacArthur, 1963], or if a concomitant cause is needed, such as species competition [Case et al., 2005, Bridle and Vines, 2007], or reducing carrying capacity [Gomulkiewicz et al., 1999].

A potential mechanism leading to the formation of sharp range margins is asymmetric gene flow from more densely populated parts of the habitat (*e*.*g*. centre) swamping locally beneficial alleles at the margins [Haldane, 1956, Mayr, 1963, Kirkpatrick and Barton, 1997, Peck et al., 1998]. However, it has been shown that asymmetric gene flow alone is insufficient to generate range margins. For example, when the optimal phenotype of the adaptive trait changes linearly in space, a population can expand indefinitely (or up to the habitat boundaries), unless the gradient in the optimum is too steep, in which case the population will face rapid global extinction or rapid fragmentation into isolated sub-populations [Barton, 2001]. In this scenario, global extinction is a deterministic event due to the mean growth rate of a population being smaller than zero. For finite populations, along gradients shallower than required for the population to get extinct deterministically, local extinction can occur because gene flow generates too much genetic variance (*i*.*e*. genetic load) – which, in turn, reduces the local population size to the point where drift overpowers selection, causing adaptation to fail everywhere [Polechová and Barton, 2015]. Conversely, if the gradient in the optimal phenotype steepens in space, adaptation will fail when the local steepness of the gradient reaches the point where too much genetic variance is needed to maintain the population at the optimum, leading to the formation of a sharp range margin [Polechová and Barton, 2015, Polechová, 2018] (see Box 1 for a more thorough overview of the framework developed by Polechová and Barton [2015]). This finding puts forward a “steepening-gradient hypothesis” for species’ range limits: the steepening-gradient hypothesis states that, in the absence of geographic barriers or competitors, range margins can form where the gradient in the optimal phenotype steepens sufficiently in space.

### Box 1

**The steepening-gradient hypothesis framework**

#### Steepening environmental gradients cause range margins

In this Box, we provide a brief summary of the seminal theoretical work on the establishment of finite ranges in the absence of geographical barriers.

The idea that the formation of range margins might be caused by gene swamping from the core of an expanding population has been proposed at least 7 decades ago by Haldane Haldane [1956]. This model was elaborated upon in a seminal paper by Kirkpatrick and Barton [1997]. Kirkpatrick and Barton [1997] considered a deterministic model of a population in a one-dimensional habitat, with one adaptive polygenic trait, such that the optimal phenotype of the trait changes linearly in space (i.e. constant environmental gradient). In addition, the genetic variance of the trait was assumed to be fixed. They showed that such a population will attain a finite range due to asymmetric gene flow between the core and the edge of the range. This is, in turn, because local maladaptation, i.e., the deviation of the mean population phenotype from the local optimal phenotype, increases from the range centre towards range margins.

However, Barton [2001] later showed that for a deterministic model with one polygenic adaptive trait, where the distribution of allelic effects at each locus underlying the adaptive trait is either Gaussian distributed or bi-allelic, and genetic variance is allowed to evolve, dispersal inflates the genetic variance rather than causing a mismatch between the mean population phenotype and the local optimum. Thus, with such distributions of allelic effects, the only stable solution is perfect adaptation everywhere, unless indefinite range expansion is prevented, and this occurs when the effective steepness of the environmental gradient (i.e., the gradient in the optimal phenotype of the adaptive trait) is so high that the growth rate becomes negative due to the genetic load caused by genetic variance. If the gradient in the optimal phenotype is constant in space, this latter solution implies rapid global extinction, as opposed to infinite range expansion. More generally, Barton [2001] advocated that, for a population to have a finite range, the optimal phenotype needs to be steepening in space – what we call the “steepening-gradient hypothesis”. A further generalization to this steepening-gradient hypothesis, particularly to include the effect of finite population size, was examined by Polechová and Barton [2015].

#### The model

In Polechová and Barton [2015]’s model, it is assumed that the population expands along a smoothly changing environmental gradient. More precisely, the individuals are assumed to have one adaptive trait under stabilizing selection towards a spatially dependent optimal phenotype, *θ*(*x*) (where *x* denotes the spatial position). The spatial rate of change *b*(*x*) of *θ*(*x*) at each point *x* (*i*.*e*. the local steepness of the gradient in the optimal phenotype) is given by *b*(*x*) = *θ*^*′*^(*x*), where *′* denotes the spatial derivative. Using simulations, Polechová and Barton [2015] found that the population undergoes range expansion as long as the **effective environmental gradient**

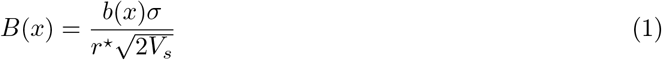

is below a certain critical threshold *B*_crit_. In eq. (1), 1*/V*_*s*_ denotes the intensity of stabilizing selection, and *r*^⋆^ stands for the growth rate of the population and is defined as *r*^⋆^ = *r*_max_ − *v*_*g*_*/*(2*V*_*s*_) (with *r*_max_ the intrinsic growth rate of the population, and *v*_*g*_ the adaptive genetic variance). In general, if *r*^⋆^ *>* 0 a perfectly adapted population will persist.

The critical effective gradient *B*_crit_ depends on the local population size *N* (*x*) and the selection coefficient per locus underlying the adaptive trait, *s* = *α*^2^*/*(2*V*_*s*_) (±*α/*2 being the additive effect of each allele on the phenotype), as well as the standard deviation of dispersal distance *σ*. That is, a local diploid population can maintain a positive population size as long as

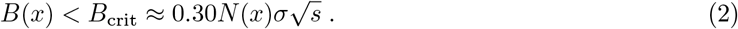

Note that the factor 0.3 appearing on the right-hand side of (2) accounts for diploidy [Eriksson and Rafajlović, 2021], whereas in haploid populations, the corresponding factor is 0.15 [Polechová and Barton, 2015].

Condition (2) implies that a population will persist locally as long as either drift is weak (that is, the local neighbourhood size *N* (*x*)*σ* is large), or the selection coefficient per locus is high. In other words, selection needs to be strong with respect to drift in order to ensure the persistence of a population for a given effective environmental gradient. Finally, note that the condition expressed in (2) was determined numerically, hence the appearance of ≈ sign on the right-hand side of (2). This means that he expression for *B*_crit_, and hence for the position of range margins, is not exact. However, simulations have shown that the population ceases to exist in the close vicinity of, although somewhat beyond, *x*_crit_ such that 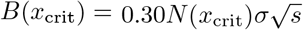 [Polechová and Barton, 2015, Eriksson and Rafajlović, 2021, 2022]. For simplicity, we often refer to the formation of “sharp margins”. However, due to dispersal, the margins that form in the kind of model developed by Polechová and Barton [2015] are not “sharp”, as the population always decreases quickly (*i*.*e*. faster than expected deterministically, see below) over a few demes. However, to the scope of this paper, we argue that this distinction is irrelevant (as explained in the Discussion; also see Eriksson and Rafajlović [2021] for a more thorough discussion of the sharpness of margins).

A crucial aspect of the Polechová and Barton [2015] model is the fact that the population size *N* (*x*) depends on the local genetic variance of the adaptive trait, *v*_*g*_(*x*). If the population is distributed along a unidimensional discrete habitat composed by demes of carrying capacity *K*, the expected population size in this model is

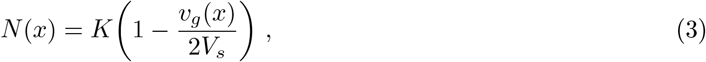

and genetic variance *v*_*g*_(*x*) is expected to increase when the steepness of the environmental gradient increases. This happens because gene flow displaces individuals that are locally adapted to places in the habitat where they are poorly adapted, thus inflating genetic variance. When the gradient is locally steeper, the average difference between the immigrants’ phenotype and the locally optimal phenotype is larger.

This box provides an insight into the key components of the steepening-gradient hypothesis. Notably, however, this hypothesis and its implications are grounded in the assumption that the environmental selection acts on a single adaptive trait. In this study we extend this by accounting for multiple (up to three) adaptive traits.

So far, the theoretical treatment of the steepening-gradient hypothesis has taken into account the evolution of a single adaptive trait. However, empirical studies have provided evidence that, in natural populations, multiple adaptive traits are likely involved (*e*.*g*. Manel and Holderegger [2013], Dayan [2020]). Here we expand the steepening-gradient hypothesis put forward by Polechová and Barton [2015] in two directions. First, we assess how the formation of range margins is impacted by the interplay between two adaptive traits under stabilizing selection towards separate optimal phenotypes, one with the gradient in the optimal phenotype that steepens in space, and one with a constant gradient (*i*.*e*. a linear optimum). Because we impose two components of selection, a lower population fitness is expected compared to when only one of the selection components (acting on a single trait) is involved [Barton, 2001]. However, because a population is expected to expand indefinitely over a linearly changing optimum, it is unclear if, and how, added selection in the form of a linearly changing optimal phenotype affects the realised range of a population. We show that individual selection gradients steepen each other, leading to smaller ranges than when any individual gradient acts alone. Notably, this can be true even when the steepening gradient – without which a finite range cannot be achieved in our modelling – is very shallow, such that it can be overlooked in field studies.

Second, we ask if, and how, range expansion is influenced when a given (composite) environmental gradient is decomposed into multiple (up to three) separate components of environmental selection, each acting on a separate adaptive trait. Here, we are specifically interested in a comparison to the situation when the same composite environmental gradient acts on a single adaptive trait. We show that a population can attain a larger range when the selection gradient is decomposed into two or three orthogonal adaptive traits than when it acts on a single adaptive trait. In addition to the range being larger, a population where stabilizing selection from a composite environmental gradient acts on multiple traits is generally less maladapted across a range compared to a population with less adaptive traits under stabilizing selection.

These results suggest increased selection for the evolution of novel traits under harsh environmental conditions.

## Materials and methods

### Model summary

Similarly to Eriksson and Rafajlović [2021, 2022], we modelled a monoecious population with non-overlapping generations expanding from a central region of a unidimensional habitat. The habitat was composed of 180 demes, each with the carrying capacity of *K* = 100 individuals (or, where stated, *K* = 200). At the beginning of the simulations, a region of 36 demes around the center of the habitat (*i*.*e*. from deme 73 to 108) was filled with locally well adapted populations (as explained in Eriksson and Rafajlović [2021, 2022]). In each generation *τ*, the fitness of individual *l* in deme *i* ∈ {1, 2, …, 180} was determined by the local population size 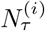, and by the individual’s phenotypes 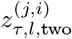 of two polygenic adaptive traits (*j* ∈ {1, 2}; the subscript ‘two’ refers to the fact that this quantity is defined for a model with two traits, opposed to models with one and three traits described below). Each trait was assumed to be under stabilizing selection towards a spatially heterogeneous optimal phenotype 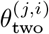, such that 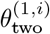 had a steepening gradient in space (eq. S1, Appendix A) and 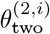 was linear in space (eq. S2, Appendix A). The diagram in figure 1 illustrates the model. Each adaptive trait was underlain by a separate set of *L* freely recombining biallelic loci. Alleles were assumed to contribute additively to the phenotype with ±*α/*2. We assumed that the two traits were orthogonal, *i*.*e*. if the trait value of one of the two traits was the same in two individuals in the same deme, then the difference in fitness of these individuals would be determined by the other trait value only. The growth rate of each individual was

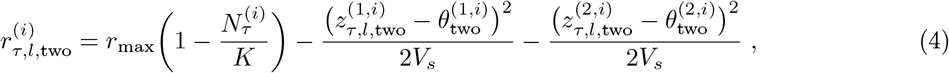

where r_max_ is the maximal intrinsic growth rate of the population (*r*_max_ = 1 throughout). The second and the third term in (4) are the contributions due to selection acting on the first and second trait, respectively, in line with Fisher’s geometric model involving two orthogonal traits [Fisher, 1930]. Here, *V*_*s*_ is the width of stabilizing selection, and we set it to the same value for both traits for simplicity (unless otherwise noted).

**Figure 1:**
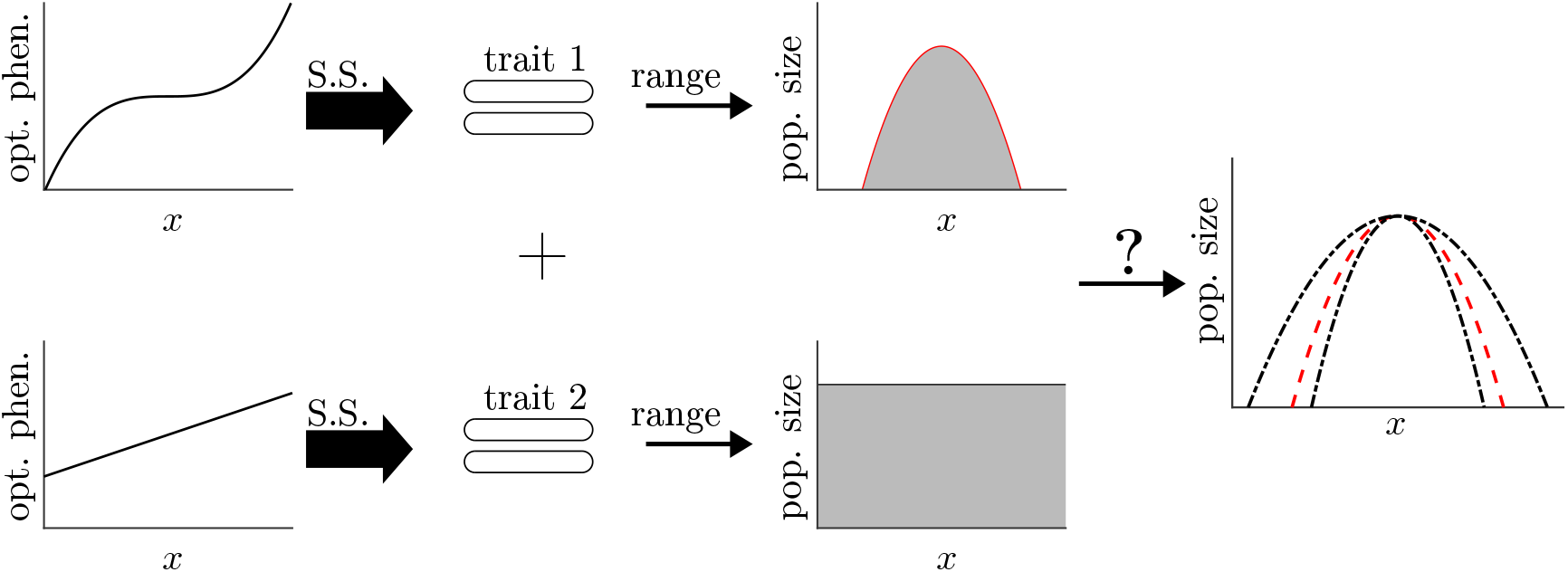
A population with an adaptive trait under stabilizing selection (S.S. in the diagram) towards an optimal phenotype steepening in space (upper panels) is expected to have a finite range [Polechová and Barton, 2015], while a population with an adaptive trait under stabilizing selection towards a linear optimal phenotype (lower panels) will have an infinite range – if the steepness of the gradient is smaller than a certain critical threshold [Barton, 2001]. In the first part of our study we consider a population with two adaptive traits, one under stabilizing selection towards a steepening optimum, the other towards a linear optimum, and we characterize the realized range with respect to the case with only the first adaptive trait and its corresponding steepening gradient in the optimal phenotype, whereas the second trait was assumed to be neutral.

To understand how an additional component of selection in the form of a constant gradient in the optimal phenotype may modify the establishment of range margins, we first compared the results obtained under this model with two adaptive traits to the results obtained under the model involving only one adaptive trait with the steepening gradient in 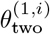. This was done by letting *V*_*s*_ → ∞ for the second trait, making it a neutrally evolving trait (see figures S1–S2). Thereafter, we aimed at understanding if a population is more or less successful in expanding its range when a composite environmental gradient is decomposed into environmental selection acting on two traits as opposed to it acting on one trait only. To achieve this, we compared the model with two adaptive traits with a model with one adaptive trait, that we dub the “composite one-trait model”, or just “one-trait model”. Note that linguistically we differentiate this from the “single-trait model”, which refers to the model outlined for a population with a single trait by Polechová and Barton [2015] (see Box 1). In the composite one-trait model, the adaptive trait was underlain by 2*L* loci with phenotype 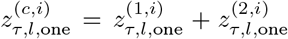 (figure 2, upper panel). Here, 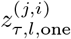 (with j ∈ {1, 2}) simply denotes *j*-th component of the phenotype that is determined by the *j*-th set of *L* loci. This splitting of the phenotype in separate components (although they contribute to the same adaptive trait) was done for convenience, and to facilitate the comparison to the two-trait model. The subscript ‘one’ in the notations above indicates that a given quantity corresponds to a one-trait model. In the composite one-trait model model, the composite optimal phenotype was 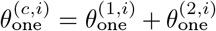, and the growth rate 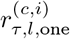 for individual *l* in deme *i* at generation *τ* was:

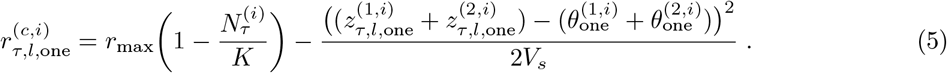

**Figure 2:**
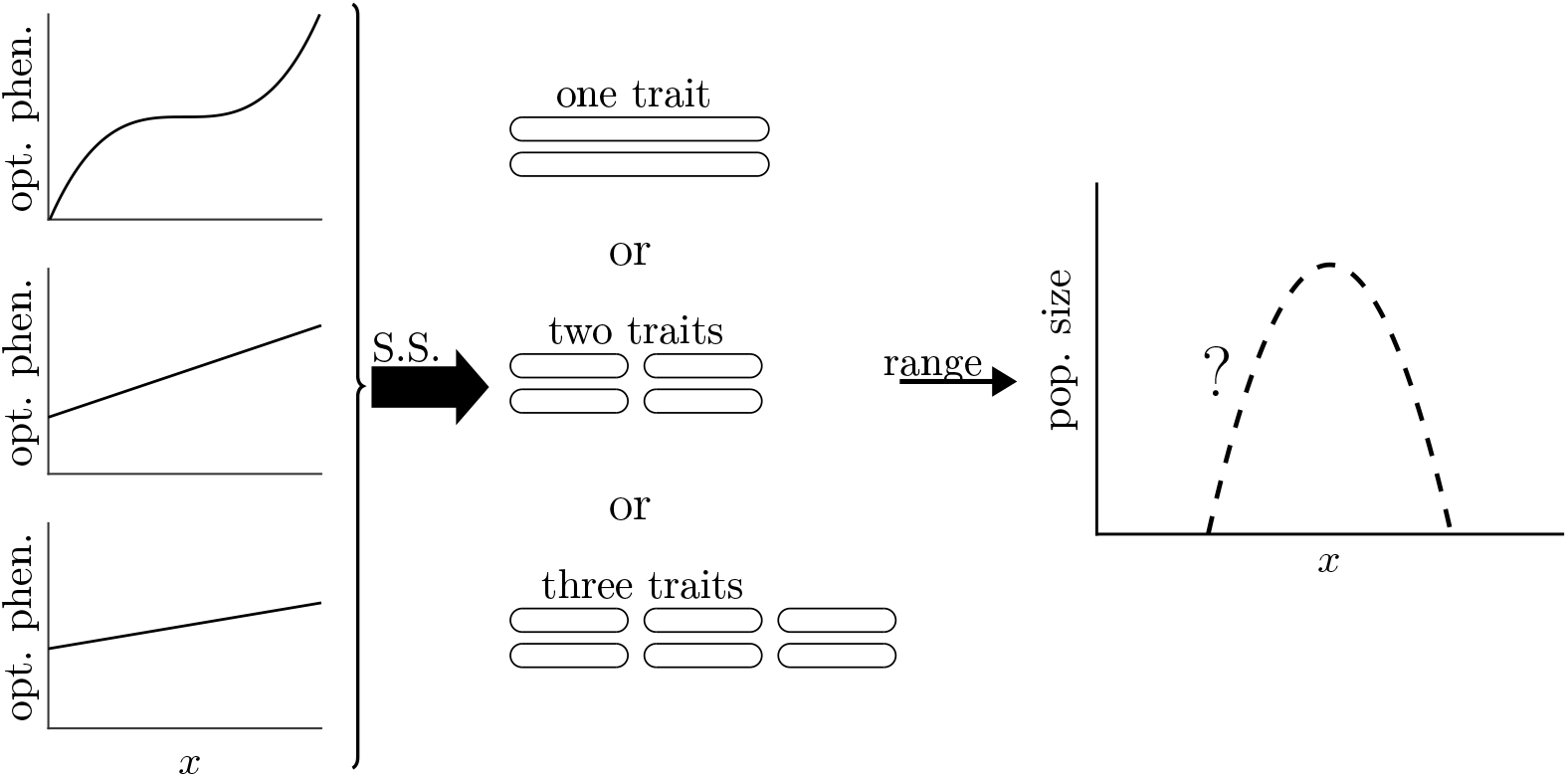
In the second part of our study, we consider a population with one, two or three traits under stabilizing selection from an environmental gradient divided in three environmental factors, the first of which has a gradient steepening in space, and the other two have a constant gradient. We aim at characterizing population size and realized range extension in the three models.

Finally, we also compared both the one-trait and the two-trait models to a three-trait model where each individual *l* in generation *τ* had phenotypes 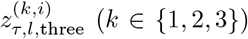 and each adaptive trait was under stabilizing selection towards a separate optimal phenotype 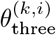 (figure 2, lower panel). For trait 1, the optimal phenotype in deme 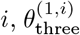, was the same as for the case with two traits. The optimal phenotypes for traits 2 and 3, 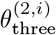 and 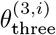, were linear with steepnesses 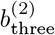 and 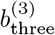 (eq. (S3), Appendix A). To compare the results of the models with one, two and three traits, we assumed that 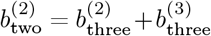. Similarly to the two-trait model, the growth rate for the three-trait model was:

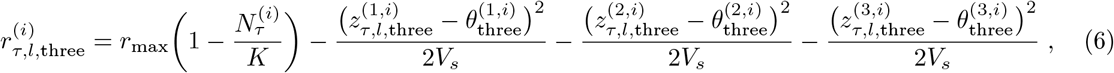

Throughout, quantities without the subscript *τ* are quantities at the end of simulations (*i*.*e*. quasi-equilibrium, *sensu* Eriksson et al. [2013], albeit, like others, *e*.*g*. Barton [2001], and for simplicity, we refer to this state as equilibrium). In the simulations, the final range was established many generations before the end of the simulation (see figure S3, Appendix A).

In each generation, after selection, recombination, mutation and fertilization took place, individuals dispersed following a discretised Gaussian distribution centered around their native deme and with standard deviation of *σ* = 1 [Eriksson and Rafajlović, 2021]. Simulations were run for 100,000 generations, with 50 independent replicates per parameter set.

In the comparison between the two-traits model and the single-trait model (*i*.*e*. with *V*_*s*_ → ∞ for the second trait), we explored 4 different values of the steepness 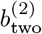 of the linearly changing optimal phenotype 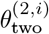. We kept 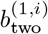 the same in each case (table 1). At the end of each simulation, before migration took place in the last simulated generation, we measured the range attained, the local population size *N* ^(*i*)^ along the range, as well as the genetic variance *v*^(*j,i*)^ for each trait. We also used the theoretical expectations (population size, genetic variance) for the single-trait model as a baseline for comparisons [Polechová and Barton, 2015, Fouqueau and Roze, 2021]. When analyzing results of the one-, two- and three-trait models, we used as a base for comparisons the case where in the two-trait model 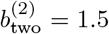. This was done for carrying capacity *K* = 100 and *K* = 200. For the three-trait model, we used 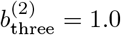 and 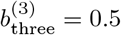.

**Table 1:**
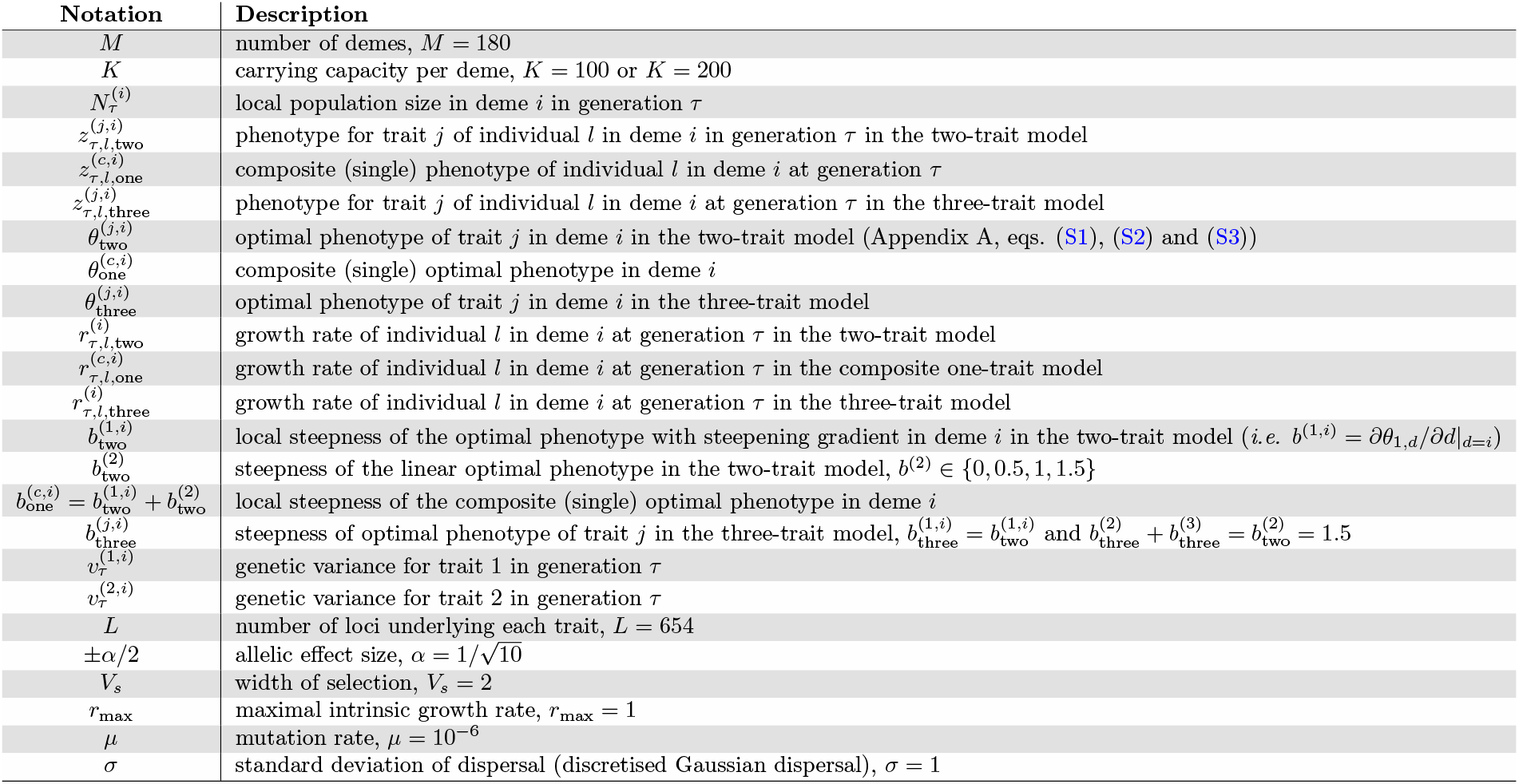
Notations used in the main text. For model parameters, the values used in our simulations are stated. For further details on model implementation and parameters, see the Appendix A.

## Results

### Adding a constant environmental gradient on a separate adaptive trait

#### Failure to adapt and reduced range

Let us start by looking at the results in the model where two adaptive traits are under stabilizing selection towards separate optimal phenotypes. First, we compare the genetic variance for each adaptive trait to the genetic variance that would be obtained if only one of the two traits was under stabilizing selection whereas the other trait evolves neutrally (the expectations for the model with only one trait are calculated in Barton [2001] and Fouqueau and Roze [2021]). Close to the centre of the habitat (that is, the demes closer to *i* = 90 and *i* = 91), the genetic variance for the trait with a spatially steepening gradient in the optimal phenotype (hereafter trait 1) follows the expected genetic variance for a population expanding over the steepening gradient alone, *i*.*e*. 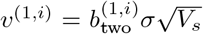 [Barton, 2001, Polechová and Barton, 2015]. However, far from the centre, this approximation over-estimates the genetic variance. This is because in the centre, the expected genetic variance is much smaller than the width of stabilizing selection *V*_*s*_ – that is, *v*^(*j,i*)^ ≪ *V*_*s*_, and the approximation presented in Barton [2001] for the genetic variance is valid when selection acting on each trait *j* is weak (*i*.*e*. when linkage disequilibrium can be neglected). Because in the cases considered here, the realized genetic variance is only slightly smaller than *V*_*s*_ away from the center and for large 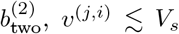, a better approximation is given by Fouqueau and Roze [2021], 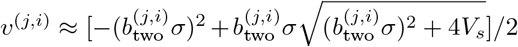 (blue dashed line in fig. 3A, also see comparison between the two formulas by Barton [2001] and Fouqueau and Roze [2021] in Appendix B, fig. S4). The genetic variance for the trait with the optimal phenotype that changes linearly across the habitat (hereafter trait 2) is approximately constant across the realised range. This is qualitatively in line with theory for a single-trait model [Barton, 2001, Fouqueau and Roze, 2021]. The spatial pattern of the variance for trait 2 is retained for all values of 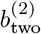 employed here, *i*.*e*. we find that the variance *v*^(2,*i*)^ is approximately spatially constant, although it is slightly concave due to the lower population size near the margins (also, realised clinal patterns were consistent with this result, see figs. S5, S6 and S7 in Appendix B).

**Figure 3:**
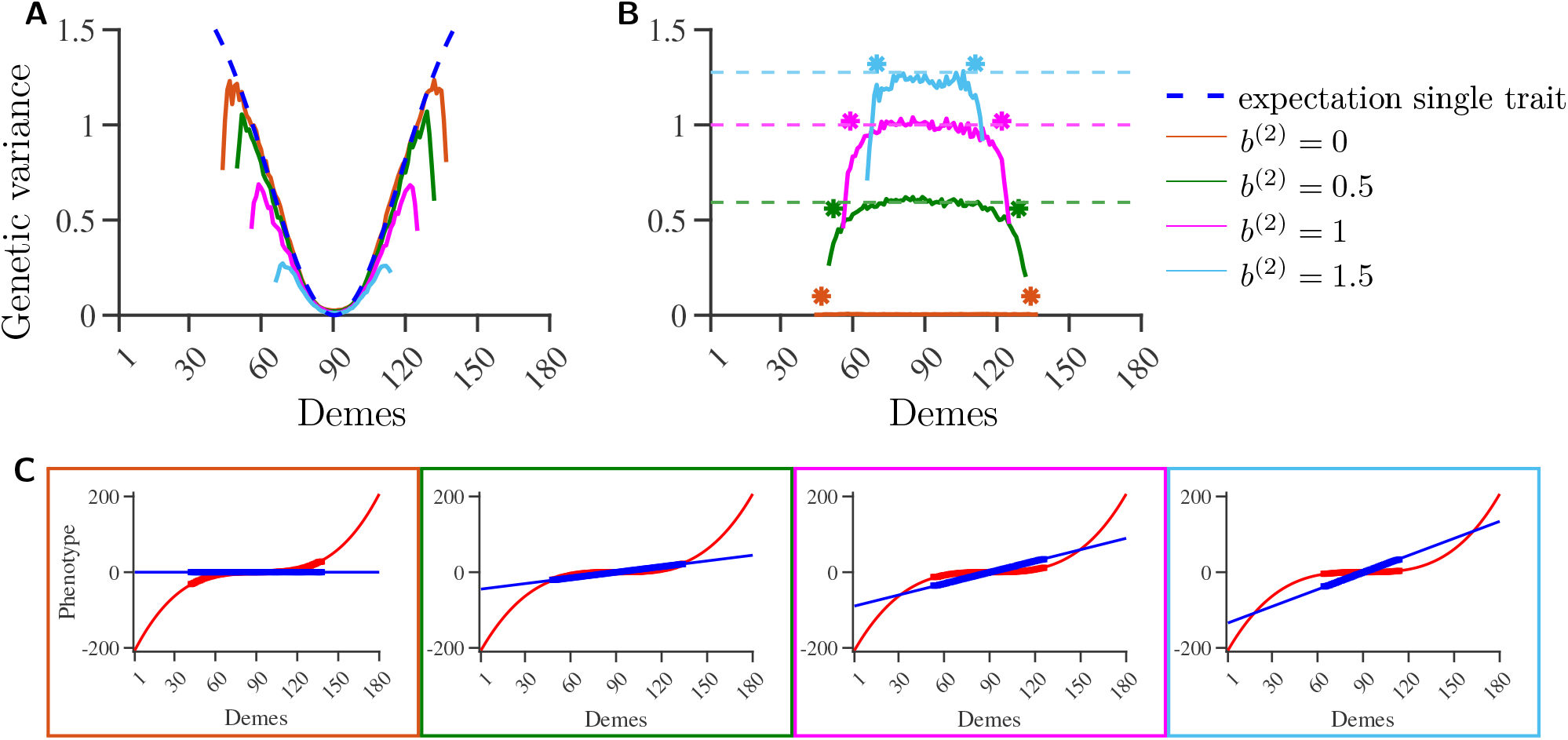
Genetic variance averaged over 50 independent realisations for the phenotype of trait 1 (A) or trait 2 (B), as a function of deme position. Results for four different values of the steepness of the constant gradient, 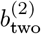, are shown (as indicated in the legend). The solid lines correspond to the two-trait model. In (A) the blue dashed line shows the expected variance for trait 1 for a single-trait model according to Fouqueau and Roze [2021], 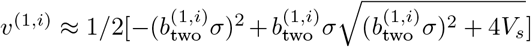. In (B), green, magenta and cyan dashed lines represent the expected variance for trait 2, 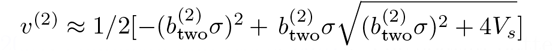 (colored according to the figure legend). The stars denote where adaptation fails according to maximum variance in (A) (colored according to the figure legend). In panel (C), we show the optimal phenotypes 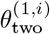 (steepening gradient; red line) and *θ*^(2,*i)*^ (constant gradient; blue line). The colors if the boxes correspond to the colors used in panels (A) and (B). Further details on the gradients are given in Appendix A. In each coloured box, the thicker line shows realised local mean phenotypes at the end of simulations, averaged over 50 independent realisations of range expansion. Other parameters: 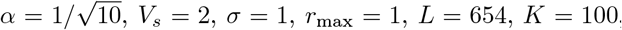, and 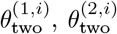 as given in Appendix A, (S1) and (S2) respectively.

We found that the realised range is smaller when 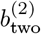 is larger (fig. 3; recall that 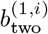 was kept the same in all runs). In other words, the two gradients steepen each other. Notably, we find that, at the margins, the steepening gradient is shallower when the second, constant, gradient 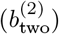 is larger. Recall that a constant gradient by itself cannot cause the formation of non-trivial range margins (*sensu* Eriksson and Rafajlović [2022]). But when a steepening gradient, despite being almost flat, is added on top of a constant gradient, a non-trivial range margin is readily formed (fig. 3C).

### Decomposing a single optimal phenotype into two or three optimal phenotypes

We aimed at understanding whether a range-expanding population attains a larger (or smaller) range when a composite environmental gradient is decomposed into environmental selection acting on multiple traits as opposed to it acting on one trait only. To this end, we compared the model with a single composite adaptive trait (individual phenotype 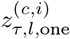, see Methods section) to the model with two orthogonal traits, and explored the same values of 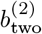 as in the previous section (fig. 5). We further explored a model with three orthogonal adaptive traits, where one trait is under stabilizing selection towards the steepening optimum, and two are under stabilizing selection towards separate linear optimal phenotypes with steepnesses 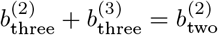.

**Figure 4:**
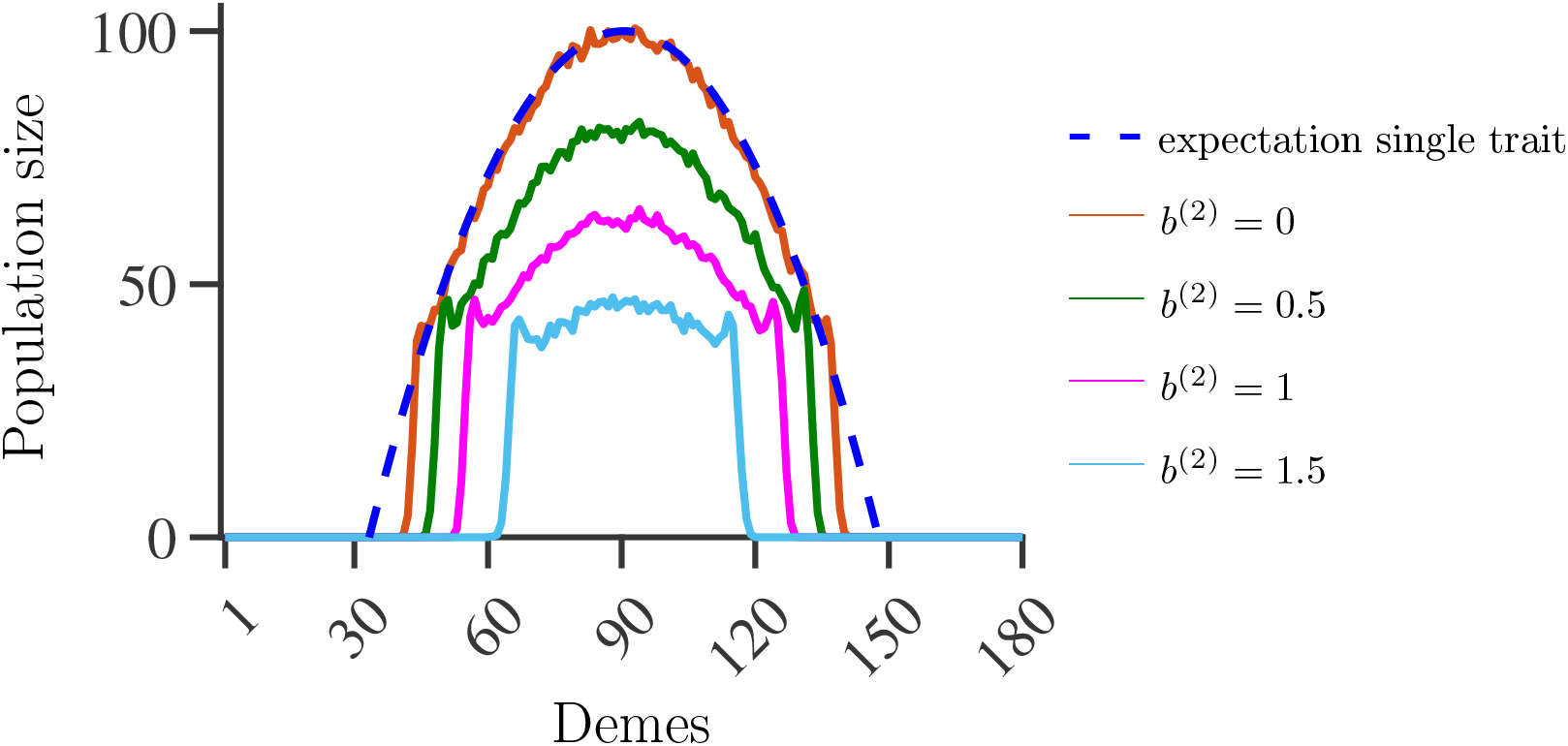
Population size averaged over 50 independent realisations for different values of the steepness of the constant gradient, *b*^(2)^, as a function of deme position. Solid lines show realised population size at the end of simulations for *b*^(2)^ = 0 (orange), *b*^(2)^ = 0.5 (green), *b*^(2)^ = 1 (magenta), and *b*^(2)^ = 1.5 (cyan). Blue dashed line shows the expected population size with only the steepening gradient *b*^(1,*i)*^, *i*.*e*. 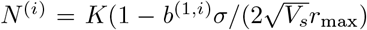. Other parameters: 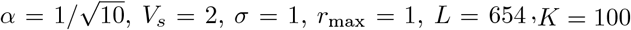, and *b*^(1)^ as defined in Appendix A (equation (S1)).

**Figure 5:**
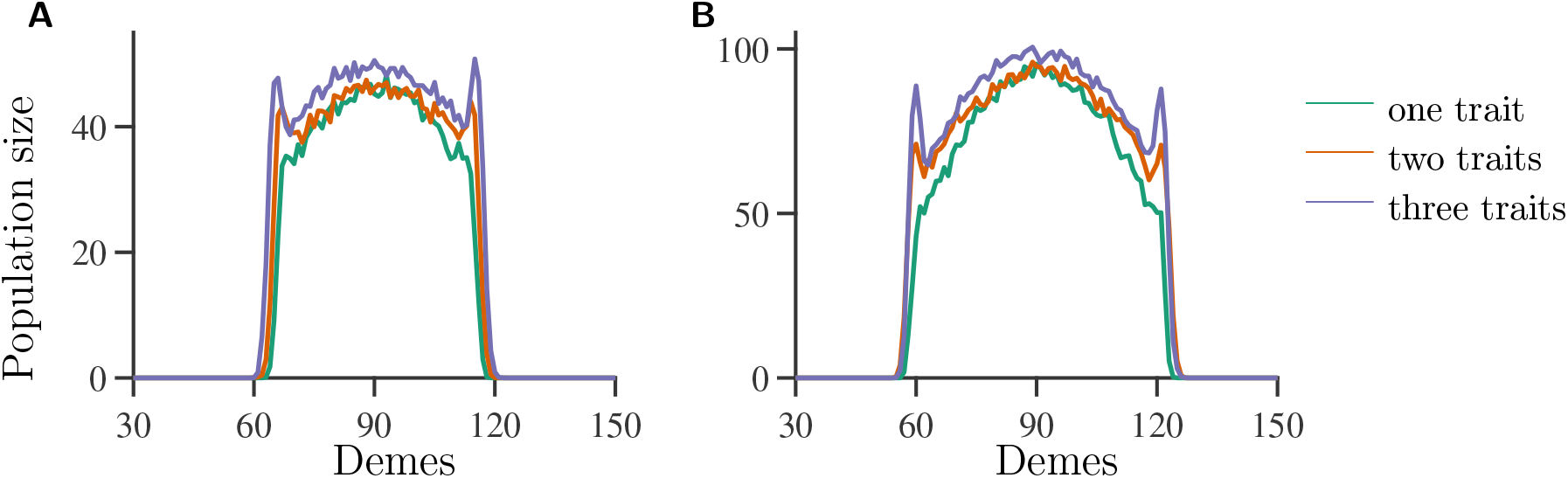
Comparison of population size between a one-trait model with single composite gradient *θ*^(*c,i)*^ (green line), a two-trait model with 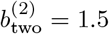 (orange line) and a three-trait model with 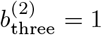 and 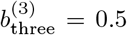 (purple line). Measurements were taken before migration and after selection. Panels (A), (B): population size averaged over 50 independent realisations for different carrying capacities, *K* = 100 (A) and *K* = 200 (B), as a function of deme position. Other parameters: 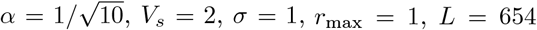, and the optimal phenotypes as given in Appendix A, eqs. (S1), (S2) and (S3) respectively.

With a single trait, the range follows closely the theoretical expectations [Polechová and Barton, 2015] with respect to the steepness of the gradient at the range margin (table S1, appendix B), as well as the expected population size (fig. 5A, B). With two or three traits, the realized range was slightly larger than in the one-trait model (figure 5). The larger realized range might in part be explained by the fact that, in general, local population size across the range was the same or larger for the two- and three-trait model than in the one-trait model. In the center, the population size in the one-trait model is comparable to the population size in the two-trait model, while at the margins a larger population size is realized when multiple traits are involved. The population size in the three-trait model was higher than in the one- and two-trait model across the whole range. Additionally, while the population size decreases monotonically from the center to the margins in the one-trait model, with multiple traits it shows a noticeable increase of population size in the margins compared to the decreasing trend. The small increase of population size at the margins is a feature of the two-trait and the three-trait model, but does not appear in the composite one-trait model. Thus going towards the margins, the population was less maladapted in the two- and three-trait model than in the single composite trait model (fig. 5A, B). This was also reflected in the mean population fitness, which was larger at the margins in the two-trait model than in the one-trait model (fig. S8, Appendix D).

## Discussion

Earlier studies employing one adaptive trait with a spatially steepening gradient in the optimal phenotype (referred to as ‘steepening environmental gradient’) showed that a population can expand over such a habitat and attain sufficient local adaptation, provided that the genetic variance increases sufficiently in space: when the local variance cannot increase sufficiently, adaptation fails and range margins form [Polechová and Barton, 2015]. By contrast, when the optimal phenotype of the adaptive trait changes linearly in space, expansion will be unlimited, or the population will face rapid global extinction or fragmentation [Barton, 2001, Polechová and Barton, 2015]. Here, we extended this single-trait model to account for two (or three) adaptive traits. We first studied the formation of range margins in a two-trait model, such that one trait has a spatially steepening and the other a spatially constant gradient in the optimal phenotype, in comparison to a model where only the first trait is adaptive and the gradient in optimal phenotype has the same shape as in the two-trait model, whereas the second trait evolves neutrally. Despite the fact that in a single-trait model, adaptation under a constant environmental gradient can proceed indefinitely (unless it results in rapid global extinction), we showed that the size of the realized range in the two-trait model is determined by the combined effect of both the steepening and the constant gradient in the optimal phenotypes (figs. 3, 4): the range is smaller in comparison to that expected in the absence of the constant environmental gradient. Consequently, a range margin may form even when the steepening gradient in the optimal phenotype is shallower than the constant gradient over the entire attainable range (fig. 3C, table S1). In fact, in the scenario with the steepest constant gradient that we simulated 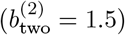, the maximum steepness of the steepening gradient over the range (realised at the margins) was approximately 4 times smaller than the steepness of the constant gradient (fig. 3C, table S1, appendix B), and yet it had a fundamental role in the establishment of the range margins.

In the second part of this study, we compared the range expansion between a population with one, two or three adaptive traits under stabilizing selection along an environmental gradient composed by one, two or three environmental factors respectively (each acting on a separate adaptive trait), while keeping the total (composite) environmental gradient the same in the model comparisons. We found that the realized range was slightly larger in the multiple- than in one-trait models. This reflects the fact that the population in the two-trait model was less maladapted across the range than in the one-trait model (except in the centre of the occupied range, where one- and two-trait models had the same extent of maladaptation, measured by the population size), while the model with three adaptive traits showed less maladaptation than the other two models across the whole occupied range. This can be understood by comparing the loss of fitness for individuals with phenotypes that deviate from the optimal phenotype. For example, for the two-trait model, we can imagine an individual with phenotypes 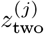 (super-/subscripts *i, l* and *τ* omitted for simplicity, *j* ∈ {1, 2} is the trait) different than the optimal phenotype, such that 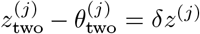.

The decrease in growth rate, Δ*r*_two_ in the two-trait model due to the deviations from optimum *δz*^(*j*)^ is (from eq. (4))

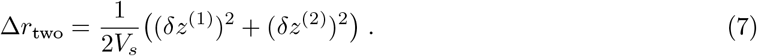

However, in the one-trait model with the optimum 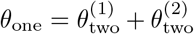, the decrease in growth rate Δ*r*_one_ as calculated from eq. (5) is

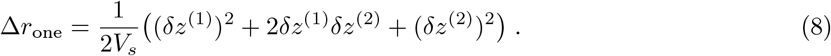

A similar calculation (not laid out here) can be performed when we take into account the fitness function for three traits under stabilizing selection towards three separate optima (eq. (6)). In this case, an additional mixed term 2*δz*^(2)^*δz*^(3)^ will arise.

Around the centre of the habitat, populations have low (almost zero) genetic variance around the optimal phenotype *θ*^(1,*i)*^, that is *δz*^(1)^ ≈ 0, and the mixed term 2*δz*^(1)^*δz*^(2)^ disappears, reducing eq. (8) to eq. (7). This also explains why the three-trait model has larger population size everywhere, as the mixed term 2*δz*^(2)^*δz*^(3)^ does not vanish in the centre of the habitat. Because genetic variance increases when going from the centre to the range margins also explains why towards the margins there is more maladaptation in the one-trait with respect to the two- and three-trait models (and, respectively, there is more maladaptation in the two- than in the three-trait model). As genetic variance spatially increases, more individuals exhibit a larger deviation from the optimal phenotype, leading to a larger difference in population sizes between the models at the range margins.

In turn, a better adaptation across the range in the two- or three-traits model allows the population to establish marginal pockets (consisting of several demes) with locally well-adapted individuals, and expand beyond the range margins attained in the one-trait model. However, genetic variance in such pockets is typically small, and this precludes range expansion beyond the pockets. Selfing could strengthen this ‘pocket’ effect, but whether or not selfing is necessary to generate such pockets is an open question (recall also that selfing occurs in the one-trait model as well, but such pockets are absent). Given that any environmental selection consists of many biotic and abiotic factors that are likely to vary in space (as well as in time), the steepening-gradient hypothesis for the formation of sharp range margins then seems rather plausible: for sharp range margins to form, it is sufficient that only one of the gradients in the environmental factors steepens in space to an extent that the total effective environmental gradient surpasses the critical value. This can occur even if the steepening gradient is range-wise much shallower than the other environmental gradients involved in adaptation, as our results here showed.

Some empirical evidence is consistent with the steepening-gradient hypothesis [Johannesson and Andre, 2006, Johannesson et al., 2020], however Kottler et al. [2021] showed that two key prediction of the steepening-gradient hypothesis, namely asymmetric gene flow and populations being more abundant in the center than in the margins, lack strong, general empirical support (see also Sagarin et al. [2006]). However, our simulations demonstrate that when the gradient in the optimum that changes linearly in space is large (large *b*^(2)^), the population size at the center of the range is not much larger than the population size close to the margins, owing to the total gradient being mostly determined by the (steep) constant gradient (cyan line in fig. 4). By further increasing the steepness of the constant gradient (but not too much, to avoid rapid global extinction) and/or by adding further components of environmental selection and respective adaptive traits, we expect that the concavity of the realised population size would decrease even further. While we still observe a decrease in population size from the centre to the margins (although we also observe pockets of larger population size at the margins, as discussed above), local conditions or different modes of density regulation could also modify population sizes in ways that our modelling does not account for. Note that the framework developed by Polechová and Barton [2015] to estimate where the range margin can arise implies the formation of sharp margins. However, due to ongoing gene flow, the realized range margins are not properly sharp, as the population size quickly reduces to zero over a few demes and not from one deme to the next. However, in the field this would be equivalent to observing transience (*sensu* Cayuela et al. [2018]) of a few individuals into a part of the environment where the establishment is impossible. For practical purposes, the margins that arise in our simulations can be thought of as sharp (also, see discussion by Eriksson and Rafajlović [2021]). We also found that the spatial patterns of the population size and fitness in the vicinity of range margins differ between the one-, the two- and the three-trait model, and that the difference between multiple traits and one trait increases as the number of traits increases. Adding such complexities and biological realism to the theoretical models (*e*.*g*. considering multiple traits and fitness trade-offs) is a promising way forward to understand the evolutionary causes to, and consequences of, limits to species’ ranges.

Throughout this study we assumed that fitness was governed by up to three adaptive traits. Although many different environmental variables can be involved in shaping the spatial patterns in optimal phenotypes of adaptive traits, one (idealised) possibility is that the multiple adaptive traits are under stabilizing selection towards the corresponding number of separate environmental variables, and that the spatial gradient in the optimal phenotype for each trait is correlated to the spatial gradient in the corresponding environmental variable. This is an ideal case for environmental association analyses, where a linear relationship between the adaptive trait value and the value of the ‘causal’ environmental variable is typically assumed (see *e*.*g*. Benito Garzón et al. [2019]). The results of environmental association analyses can then be used to guide management and conservation actions. For example, “assisted migration” can be implemented to achieve an estimated “trait shift” needed to avoid severe maladaptation (and hence extinction) under future environmental conditions, or to simply assist the movement of a population towards new areas where the population would suffer from a smaller (estimated) risk of maladaptation (see *e*.*g*. Rellstab et al. [2021]). Notably, our results show that situations exist such that a steepening gradient in the optimum of an adaptive trait can be very shallow (almost flat) throughout the entire contemporary range (but not necessarily outside of the realised range), and yet it can have a fundamental role in the establishment of range margins. In such situations, both the causal steepening environmental gradient and the adaptive trait under the corresponding environmental selection can be missed or overlooked (due to potential statistical power issues associated with a small steepness of the gradient), and this can have serious negative consequences on assisted migration and similar management and conservation actions. It proved difficult to provide a general mathematical extension to generalize the effective genetic variance contributing to local population size for two and three traits. Despite the mathematical complexity of the problem, our analysis still suggests that the formation of range margins generally depends on the net genetic load stemming from all adaptive traits involved – that is, different environmental factors combine to form one composite effective environmental gradient, which will determine the population distribution along a transect. Landscape [Manel and Holderegger, 2013] and seascape genomic studies [Dayan, 2020] suggest that many environmental variables may be involved in shaping spatio-temporal population dynamics. If all the underlying environmental variables had constant gradients in their corresponding optimal phenotypes, this would correspond to a scenario in which a population evolved along one composite effectively constant gradient. As long as the composite optimal phenotype changes linearly across the habitat, a population is expected to adapt indefinitely – provided that it does not encounter “trivial” margins (*sensu* Eriksson and Rafajlović [2022]). However, if any of the gradients steepens at some point in space, this can potentially lead to the formation of a range margin. When the composite effective gradient is steeper (*i*.*e*. there are more components of environmental selection that may or may not act on separate traits), the change needed in only one of the optimal phenotypes to potentially form a range margin becomes smaller. This reinforces the importance of the steepening-gradient hypothesis: in a scenario involving multiple environmental gradients acting on two or more adaptive traits, a small change in the local steepness of the gradient of only one optimal phenotype is enough to cause the establishment of a sharp range margin.

Previous works have addressed the role of multivariate selection in favoring or constraining local adaptation [Blows and Hoffmann, 2005, Blows, 2007, Agrawal and Stinchcombe, 2009, Kirkpatrick, 2009, Duputié et al., 2012, White and Butlin, 2021]. Most of these have focused on the role that genetic correlations between multiple traits have in constraining adaptation (*e*.*g*. [Agrawal and Stinchcombe, 2009, Kirkpatrick, 2009]). The role of multivariate selection on species’ ranges was explored by Duputié et al. [2012] in a scenario where a population experiences multivariate environmental gradients shifting in time. In such a scenario, no qualitative differences were found comparing the multivariate model to a univariate model such as the one in Pease et al. [1989] or Kirkpatrick and Barton [1997]. However, none of these previous works quantitatively compared the change in species’ range between a model with a single adaptive trait and multiple adaptive traits. In the approach that we took of comparing the realized range in a model with univariate or multivariate selection, we found quantitative differences, suggesting that multivariate selection can indeed favor expansion and adaptation along an environmental gradient. In our model, we made several simplifying assumptions, such as free recombination, and absence of plasticity. When recombination between loci is reduced, the steepening-gradient hypothesis is still valid [Eriksson and Rafajlović, 2021]. Reduced recombination between adaptive loci may (at least transiently) mitigate the effects of migration load [Eriksson and Rafajlović, 2021]. Similarly, we may expect transiently larger range extents in our model if the loci of one or both of the traits recombined less frequently. However, it is an open question how the recombination rate between loci underlying separate adaptive traits would impact on the rate of the range expansion and the establishment of range margins.

When it comes to phenotypic plasticity, theoretical analysis showed that, while expanding along a steepening gradient, if plasticity comes at no, or very low cost, the formation of a range limit may be suppressed [Eriksson and Rafajlović, 2022]. However, if plasticity is sufficiently costly, it can increase the range extent, but not indefinitely [Eriksson and Rafajlović, 2022]. In such cases the steepening-gradient hypothesis is still valid. While empirical evidence for plasticity costs is sparse [Auld et al., 2010, Murren et al., 2015], it has been argued that the presence of range margins along smooth gradients (not mediated by other factors) is indirect evidence that costs and/or limits to plasticity exist [DeWitt et al., 1998].

In sum, our study shows that, in the absence of any obvious geographical barriers, to obtain a finite range it is sufficient that at least one of multiple traits is under stabilizing selection towards a steepening effective environmental gradient. Note that the effective environmental gradient (defined in equation (1)) depends also on dispersal range and selective coefficient, as well as the intrinsic growth rate of a population. However, in our model we assumed that dispersal range, growth rate and selective forces at play for each trait were homogeneous across space. Under such conditions, for range margins to form it would be enough that one of the optimal phenotypes is steepening. When the number of environmental factors with a spatial gradient acting on different adaptive traits is larger, the required change in just one of the gradients, that is sufficient to cause range margins, is smaller. Such shallow gradients can be easily overlooked in field studies. We further showed that if the selection pressure from the environment is “decomposed” between multiple traits (meaning that separate traits are under stabilizing selection towards separate optima), the population can withstand higher genetic load, which in turns means that, before adaptation fails abruptly, populations can evolve more genetic variance and expand further when selection is decomposed between multiple adaptive traits, than when it acts on a single adaptive trait. Because of this, the steepening-gradient hypothesis may be consistent with the evolution of novel traits that might help a population to expand into a new territory, an observation that has been highlighted *e*.*g*. in Santos et al. [2017]. In addition, climate change has recently been linked to rapid trait evolution (*e*.*g*. Mackin et al. [2021]).

We argue that our results support the steepening-gradient hypothesis as an innate explanation of species’ range limits (when geographical barriers are absent), without the need for extensively steepening gradients in any individual trait optimum.

## Acknowledgments

We thank Roger K. Butlin for discussions on the topic and insightful comments on earlier versions of the manuscript, and members of the Linnaeus Centre for Marine Evolutionary Biology (CeMEB, https://www.gu.se/en/cemeb-marine-evolutionary-biology) for feedback and discussions on this topic. This work was funded by the Swedish Research Council Formas to KJ and MR (grant number 2019-00882). MR was also supported by a grant from Vetenskapsrådet (to MR; grant number 2021-05243), Hasselblad Foundation Grant for Female Scientists (to MR), and by the European Research Council (through CeMEB). Simulations were enabled by resources provided by the Swedish National Infrastructure for Computing (SNIC) at the Uppsala Multidisciplinary Center for Advanced Computational Science (UPPMAX, project SNIC 2021-22-286), and at the High Performance Computing Center North (HPC2N, project SNIC 2021-5-501), both partially funded by the Swedish Research Council through grant agreement no. 2018-05973.

## Authors contributions

MR conceived the study. ME and MR designed the simulations with input from MT. ME wrote the code of the simulations. MT ran simulations. MT analyzed simulation results with input from ME and MR. MT did the theoretical calculations with input and feedback from ME and MR. MT, ME and MR interpreted the results with input from KJ. MT wrote the first draft of the manuscript. All the authors revised and edited the manuscript. MR supervised the study.

## Conflicts of interest

The authors have no conflict of interests to declare.

## Availability of data

The data that support the findings of this study are on Dryad as a private repository. The repository will be made publicly available upon publication. The DOI is communicated privately to the editors to ensure double-blind conditions.

## Notes

### Competing Interest Statement

The authors have declared no competing interest.

### Summary of Updates

Updated following external review. In particular, analysis of model with three traits was added to give a more complete picture. Additionally, a box explaining the base theoretical results upon which our work is done is now present.

