## Appendices and supplemental figures for "Shallow environmental gradients can cause range margins to form"

October 28, 2022

#### Appendix A: Supplementary modelling details

##### Two-trait model

In this appendix, we provide further details about how we modelled optimal phenotypes along the habitat (fig. 3 in main text). We modelled a habitat with $M = 180$ demes. For the trait with a steepening gradient, the optimal phenotype $\theta^{(1,i)}$ in deme $i$ was given by a polynomial of third degree:

$$\theta^{(1,i)} = 2.88305743\times10^{-4}\cdot(i-1)^3 - 7.7410092\times10^{-2}\cdot(i-1)^2 + 6.92820323\cdot(i-1) - 206.691396369885\ . \quad \text{(S1)}$$

The coefficients are calculated in Matlab using the `polyfit()` function, fitting a polynomial of the second degree for the steepness of $\theta^{(1,i)}$ such that $\partial\theta^{(1,i)}/\partial i|_{i=1} = 4\sqrt{3}$ (and the same slope for deme 180; the steepness at the first and last deme was chosen in order to ensure the formation of a margin within the limits of the habitat), and steepness 0 between deme 90 and 91. We then integrated the formula obtained for the steepness, and chose the fourth coefficient of equation (S1) such that at the center of the habitat (denoted by $i^\star$, realised here between demes 90 and 91), $\theta^{(1,i^\star)} = 0$ as well as $\partial\theta^{(1,i)}/\partial i|_{i=i^\star} = 0$ (for further details, see code on Dryad: doi:10.5061/dryad.4b8gthtfg).

For the trait with a constant gradient in the optimum, we assumed that the optimal phenotype $\theta^{(2,i)}$ in deme $i$ is a linear function with slope $b^{(2)}$, such that $\theta^{(2,i^\star)} = 0$ for $i^\star$ in the centre of the habitat, that is:

$$\theta^{(2,i)} = b^{(2)} \cdot (i - 90.5)\ . \quad \text{(S2)}$$

Note that optimal phenotypes $\theta^{(j,i)}$ are defined as continuous functions but are evaluated at the discrete points $i \in \{1, 2, \ldots, M\}$.

##### Three-trait model

The optimal phenotype for trait 1 is the same for all models with multiple traits, *i.e.* $\theta^{(1,i)}_{\text{three}} = \theta^{(1,i)}_{\text{two}}$ as defined in eq. (S1). For the the two optimal phenotypes with constant gradient, these are defined in the same way as for the two-trait model (eq. (S2)):

$$\theta^{(j,i)}_{\text{three}} = b^{(j)}_{\text{three}} \cdot (i - 90.5)\ , \qquad j \in \{2, 3\}\ . \quad \text{(S3)}$$

#### Single trait simulations

To further test the validity of our simulations with two traits, we reduced the model with two *adaptive* traits to the model with one *adaptive* trait (for which theory is readily available [Barton, 2001, Polechová and Barton, 2015]), by performing simulations with one of the two traits with infinite width of stabilizing selection. With this setting, the trait with $V_s \to \infty$ was evolving neutrally. We also used the simulation results obtained under this single-trait model to compare against the results obtained under the model with two adaptive traits to understand how the interplay between two adaptive traits affects range formation during range expansions.

In figure S1, $V_s \to \infty$ for the optimal phenotype with a steepening gradient (corresponding to trait 1), while in figure S2, $V_s \to \infty$ for the optimal phenotype with constant gradient (corresponding to trait 2). In practice, when simulating this in Matlab, $V_s$ is set to `Inf`, so that $1/(2V_s)$ is evaluated as zero. The genetic variance of the trait evolving neutrally tends to the expectation for neutral genetic diversity, $v_g = HL\alpha^2$ with $H$ denoting the total heterozygosity (calculated as $H = 4N_{\text{tot}}\mu/(1 + 8N_{\text{tot}}\mu)$), $L$ the number of loci and $\alpha/2$ the allelic effect (fig. S1D and S2C).

When only trait 1 is under selection (fig. S1), average genetic diversity is in accordance with the expectation (eq. (S4)). When only trait 2 is under selection (fig. S2), the population expands to the limits of the habitat, as expected for a constant gradient, and the deviation from the expected value of genetic variance is in line with what is observed in the case when both traits are under selection (orange line in figure S2D, showing the realised genetic variance for trait 2 in the two-trait model). This confirms that the discrepancy between the expected and realised genetic variance for trait 2 in the two-trait model is not due to the interplay between two adaptive traits. The discrepancy is likely due to linkage disequilibrium inflating genetic variance more for steeper gradients and stronger drift [Felsenstein, 1977]. This further suggests that genetic variances of individual traits in our two-trait model evolve largely independently of each other (but see also next subsections).

In figure S2E, the population size is inflated at the margins of the habitat because we assumed habitat boundaries to be reflective. This means that, the edge populations only receive immigrants from one direction, *i.e.* for deme 1 (180), all immigrants come from populations that have larger (smaller) optimum than the focal deme, as opposed to all other populations that receive immigrants from both directions. In addition, the cumulative number of immigrants is smaller for edge demes than for any other deme. These two effects contribute to lowering the genetic variance (migration load) in the edge demes, allowing them to acquire larger average local population fitness and consequently to establish a larger local population size.

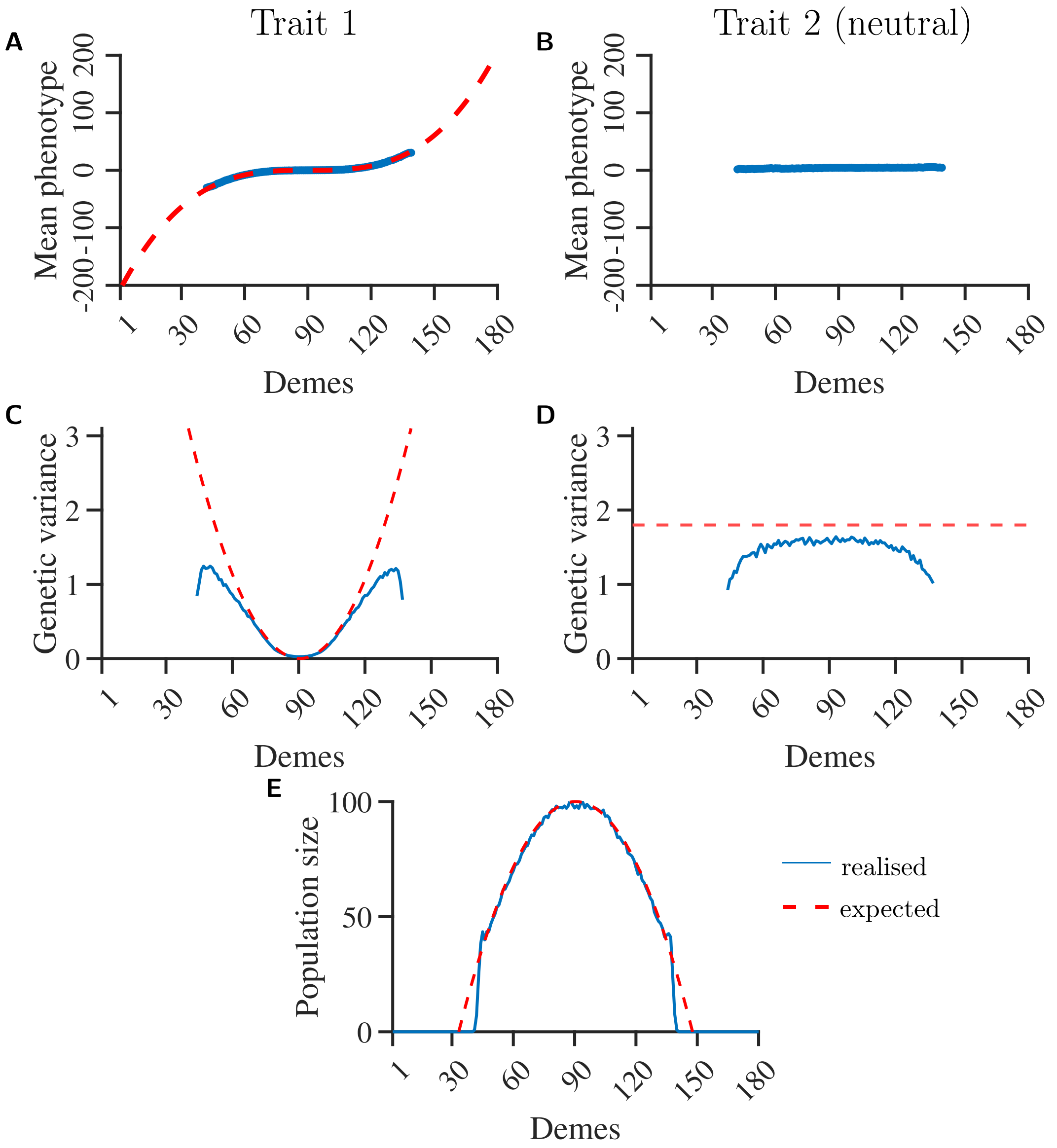


Figure S1: Range expansion in a model with a single adaptive trait that has a steepening gradient in the optimal phenotype. Trait 2 here has $V_s = \infty$. Upper row shows mean (blue dots) and expected (dashed red line) phenotype for (A) trait 1 and (B) the neutral trait 2. Middle row shows realised genetic variance for (C) trait 1 and (D) trait 2, with expected variance according to Barton [2001]. Figure (E) shows the realised population size with only trait 1 being adaptive. Adaptation fails abruptly at demes 47 and 134, consistent with equation (2) in the main text, as well as the results for the single-trait model in Eriksson and Rafajlović [2021]. Other parameters: $\alpha = 1/\sqrt{10}$, $\sigma = 1$, $r_{\max} = 1$, $L = 654$.

#### Range in equilibrium

After a few thousand generations, the expanding population reaches a state of quasi-equilibrium. As described in the Methods section, this state is not a proper equilibrium, because global extinction will

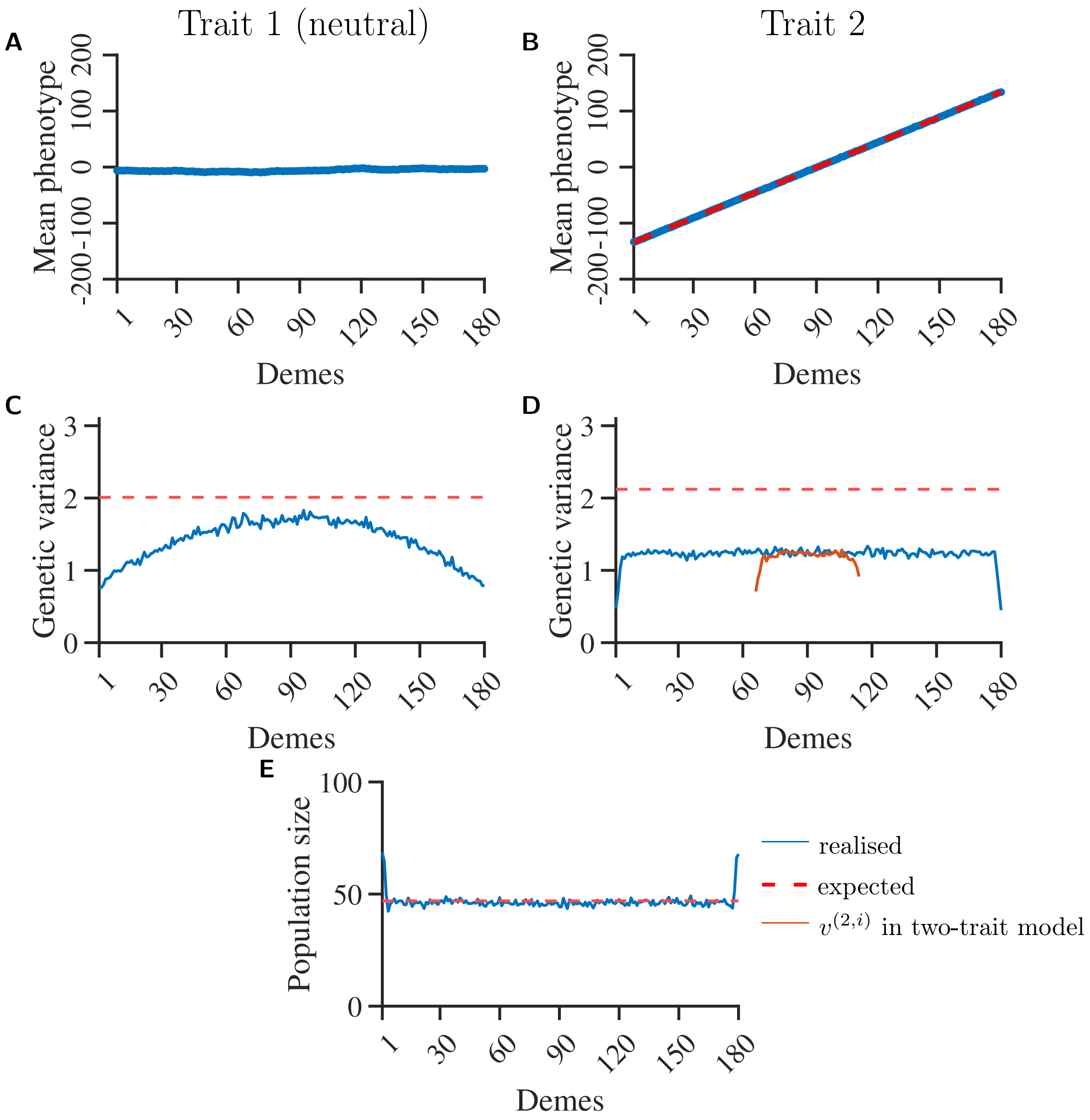


Figure S2: Range expansion in a model with a single adaptive trait that has a constant gradient in the optimal phenotype. Trait 1 here has $V_s = \infty$. Upper row shows mean (blue dots) and expected (dashed red line) phenotype for (A) the neutral trait 1 and (B) trait 2. Middle row shows realised genetic variance for (C) trait 1 and (D) trait 2, with expectations according Barton [2001]. For comparison, the orange line in panel D shows the realised variance for trait 2 when both traits are under selection (*i.e.* when $V_s = 2$ also for trait 1, data as in fig. 3). Figure (E) shows the realised population size when only trait 2 is under selection. Population size in the first and last deme is larger than expected because of reflecting boundaries. Other parameters: $\alpha = 1/\sqrt{10}$, $\sigma = 1$, $r_{\max} = 1$, $L = 654$.

ultimately occur, *albeit* this is expected after much longer times than those simulated here [Eriksson et al., 2013]. We expect the quasi-equilibrium to be long lasting – at least in terms of timescales that are of biological interest – and we confirm that this is the case in simulations (fig. S3). For simplicity, hereafter (as well as in the main text) we refer to this state as *equilibrium*.

In our model, the initially occupied range is between deme 73 and 108. For high $b^{(2)}$ (magenta and cyan

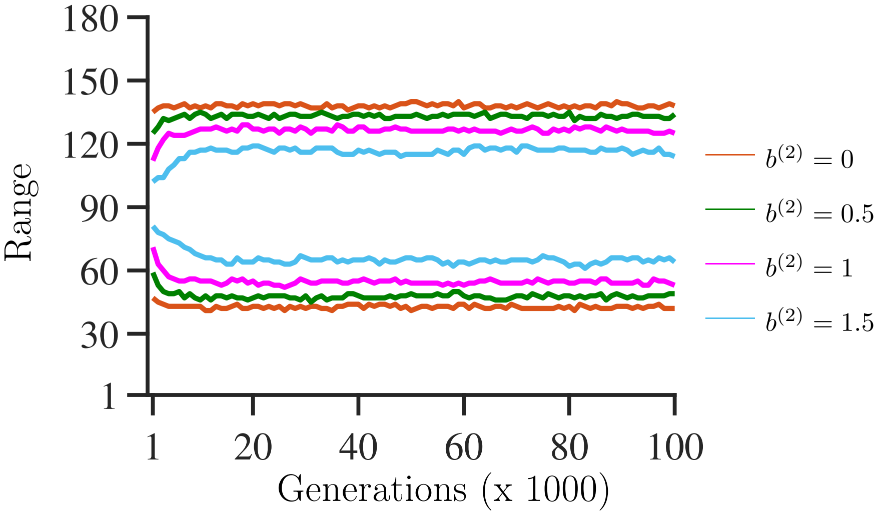


Figure S3: Range extension in a randomly chosen replicate, as a function of time for values of $b^{(2)} \in \{0, 0.5, 1, 1.5\}$. The range for $b^{(2)} = 0$ is the same as for the model with the steepening gradient only. The range here is determined as the distance between the left- and rightmost deme where $N^{(i)} > 10$, 10 being an arbitrary value. Measurements are taken every 1000 generations (first measurement at $\tau = 1000$). Other parameters: $\alpha = 1/\sqrt{10}$, $V_s = 2$, $\sigma = 1$, $r_{\max} = 1$, $\theta_{\text{two}}^{(1,i)}$ as defined in formula (S1).

lines in fig. S3), the range initially contracts in part because of slight deviations in the initial conditions due to rounding off when assigning the starting allele frequencies. However, harsh environmental conditions (steep gradients) could potentially lead to ranges smaller than initial ranges. In all the cases considered here, the range in equilibrium was larger than the initial range.

### Appendix B: Genetic variance in equilibrium and clinal patterns

#### Comparison of Barton [2001] VS Fouqueau and Roze [2021]

Barton [2001] studied the expansion of a population over a continuous unidimensional space with a spatially linearly changing optimal phenotype (*i.e.* a constant gradient). In particular, Barton [2001] found that for a trait under stabilizing selection tracking a gradient in the optimal phenotype, the spatial pattern of allele frequencies at adaptive loci in equilibrium corresponds to a series of staggered clines. For a population to track a gradient in the optimal phenotype with local steepness $b$, it is necessary to have $b/(2\alpha)$ clines per unit of distance. Each cline contributes to total genetic variance, such that the total trait genetic variance in equilibrium ($v$) depends on the slope $b$ of the optimal phenotype, as

$$v = b\sigma\sqrt{V_s}\ , \tag{S4}$$

with $\sigma$ being the standard deviation of the migration distance, and $V_s$ the width of stabilizing selection. For an optimal phenotype that changes non-linearly in space, $\theta(x)$, the local genetic variance in equilibrium is $v(x) = b(x)\sigma\sqrt{V_s}$. Here, $b(x)$ is the local slope at point $x$, determined as $b(x) = \partial b(x)/\partial x$. The genetic variance in equilibrium can be obtained from solving the differential equations describing the spatial patterns of allele frequencies at each locus [Barton, 2001]. Eq. (S4) is valid for weak selection, that is when $v \ll V_s$. In our work, however, we observe that the genetic variance $v^{(j,i)}$ is often close to the value of $V_s$ (see *e.g.* fig. 3). In the case of strong selection, the genetic varianec genetic variance $v_g$ for a single trait under stabilizing selection towards an optimal phenotype with a spatial gradient of steepness $b$ at deme $i$ before migration is [Fouqueau and Roze, 2021]

$$v_g \approx \frac{1}{2}[-(b\sigma)^2 + b\sigma\sqrt{(b\sigma)^2 + 4V_s}]\ . \tag{S5}$$

This equation approximates well the genetic variance $v^{(j,i)}$ for each trait in our model, with $b = b^{(j,i)}$. We compare the genetic variance for trait 1 realized in our two-trait model to the formulas by Barton [2001] and Fouqueau and Roze [2021] in fig. S4.

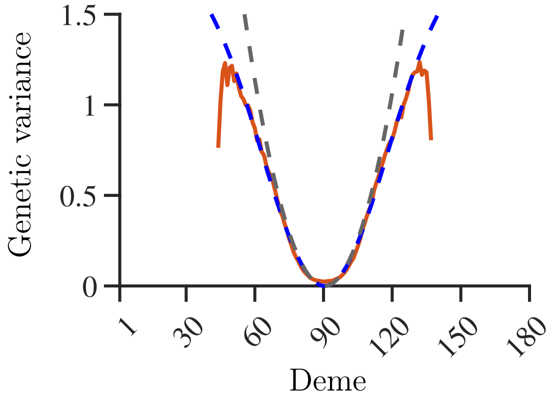


Figure S4: Realized genetic variance for trait 1 with $b^{(2)} = 0.0$ (orange solid line; same data as in figure 3) compared to the expected genetic variance in Barton [2001] (gray dashed line) and Fouqueau and Roze [2021] (blue dashed line).

#### Clinal patterns

As mentioned in the previous sections, Barton [2001] showed that allele frequencies at adaptive loci underlying trait $j$ are a series of staggered clines (fig. S5), spaced $2\alpha/b^{(j)}$ and with expected width $w = 4\sigma\sqrt{V_s}/\alpha$. In our simulation results, the realised cline spacing in equilibrium is in good agreement

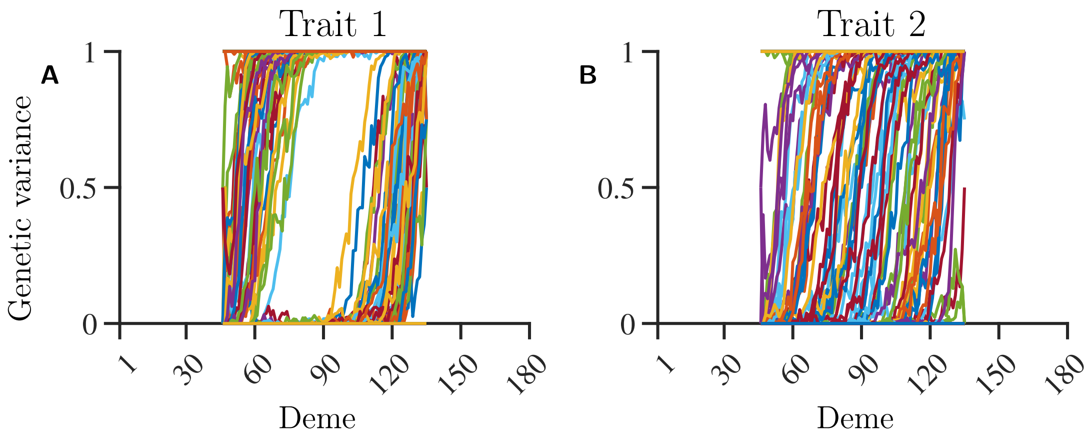


Figure S5: Allele frequency clines for a randomly chosen realisation of the two-trait model (with $b^{(2)} = 0.5$). Panel (A) shows clines for loci underlying trait 1; panel (B) shows clines for loci underlying trait 2. Other parameters: $\alpha = 1/\sqrt{10}$, $V_s = 2$, $\sigma = 1$, $r_{\max} = 1$, $\theta^{(1,i)}$ as defined in formula (S1).

with this theoretical result, but some deviations occur, particularly in the central regions where the clines obtained in the simulations are more dense than expected under a deterministic approximation (fig. S6). Because in our simulation results, the realised LE component of the genetic variance for both traits is lower than expected under the deterministic theory, the clines realised in the simulations are also narrower than expected.

The width of the clines is not uniformly distributed along the realised range. In particular, the clines

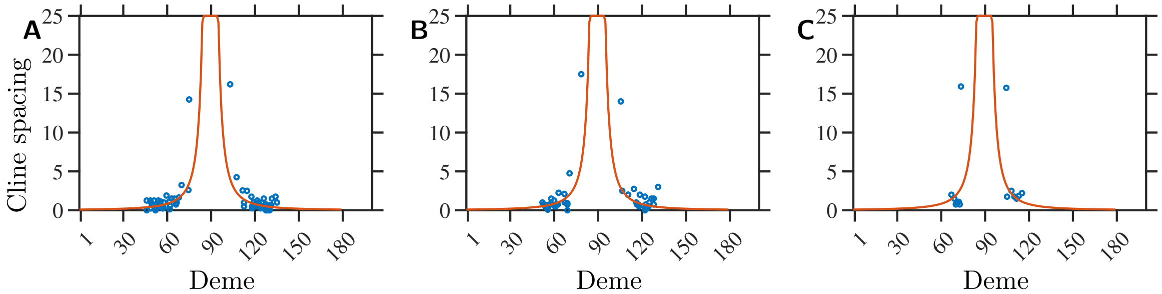


Figure S6: Spacing between the centers of the allele frequency clines for trait 1 in a randomly chosen realisation, as a function of deme number, and depending on the steepness of the linear optimal phenotype: (A) $b^{(2)} = 0.5$; (B) $b^{(2)} = 1$; (C) $b^{(2)} = 1.5$. Red line is the expected cline spacing that depends on the steepness $b^{(1,i)}$ (the spatial pattern of $b^{(1,i)}$ was kept the same in all simulated cases). Other parameters: $\alpha = 1/\sqrt{10}$, $V_s = 2$, $\sigma = 1$, $r_{\max} = 1$, $\theta^{(1,i)}$ as defined in formula (S1).

with centres closer to the range margins are narrower (figure S7) as was also find by Polechová and Barton [2015], Eriksson and Rafajlović [2021] in a single-trait model. This is, in part, due to the larger discrepancy between the LE part of the genetic variance and the expected variance in the range margins (compared to the center), which is partly due to stronger drift closer to the margins. In addition, stronger drift *per se* tends to make clines narrower [Polechová and Barton, 2011].

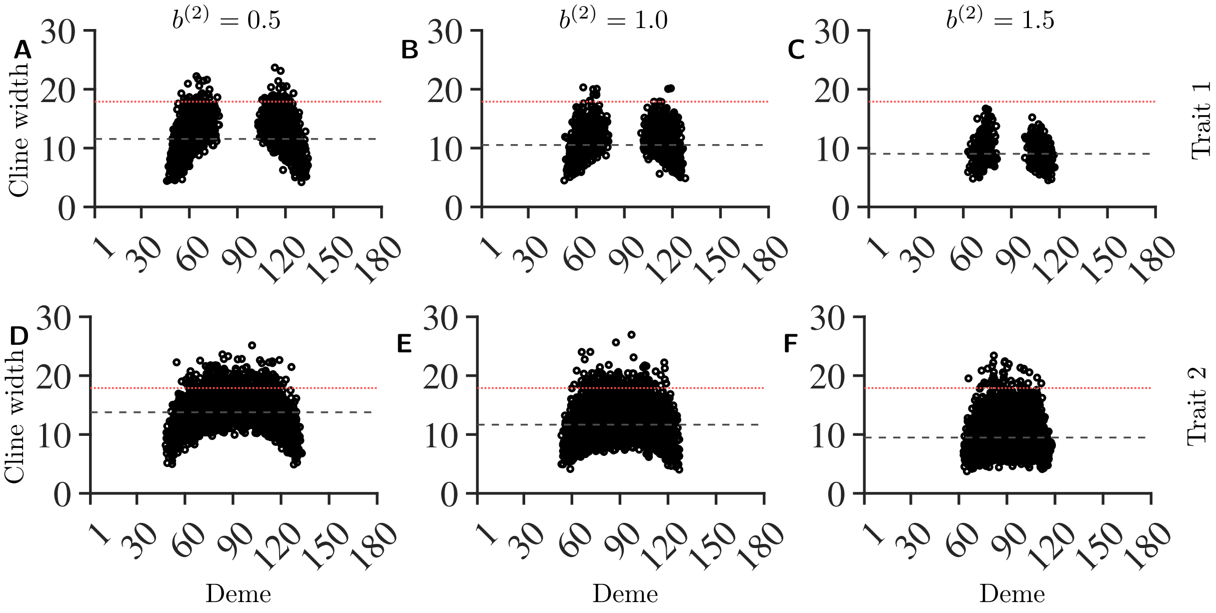


Figure S7: Cline width as a function of the cline center, for all 50 replicates for trait 1 (top row) and trait 2 (bottom row), for different steepness of the constant gradient: $b^{(2)} = 0.5$ (A), (D); $b^{(2)} = 1$ (B), (E); $b^{(2)} = 1.5$ (C), (F). The horizontal red line shows the expected cline width, while the dashed black line shows the mean realised width over all replicates. Expected width in the absence of drift is $4\sigma\sqrt{V_s}/\alpha \approx 17.9$. Other parameters: $\alpha = 1/\sqrt{10}$, $V_s = 2$, $\sigma = 1$, $r_{\max} = 1$, $\theta^{(1,i)}$ as defined in formula (S1).

### Appendix C: Differences between two-trait and composite single-trait model

In the main text, we set to understand whether a range-expanding population attains a larger (or smaller) range when a composite environmental gradient is decomposed into environmental selection acting on two traits as opposed to it acting on one trait only. In the composite single-trait model, two different sets of loci contributed to the same phenotype $z^{(c,i)}$, with the optimal phenotype defined as $\theta^{(c,i)} = \theta^{(1,i)} + \theta^{(2,i)}$. By writing, for convenience, the composite phenotype as the sum of the phenotype contribution of each set of loci, $z^{(c,i)} = z^{(1,i)} + z^{(2,i)}$, we can compare this model to the two-trait model with respect to the quantities of interest, as explained next.

##### Realised range

The range margin obtained in equilibrium, defined as the point beyond which adaptation fails (as steepness becomes too high), is larger in the two-trait model than in the single-trait model, as shown in table S2. Note that in the single-trait model, for $b^{(2)} = 1.5$, genetic variance was approximately constant over the realised range, making it difficult to estimate where adaptation fails. The value reported in table S2 (and composite steepness reported in table S1) corresponds to the numerical maximal genetic variance, but visual inspection suggests that the range calculated in table S2 is smaller than the realised critical range.

| steepness $b^{(2)}$ | $b^{(1,m)} + b^{(2)}$ [mean (SD)] | |
|---|---|---|
| | two-trait model | single-trait model |
| $b^{(2)} = 0$ | 1.6 (0.3) | 1.6 (0.3) |
| $b^{(2)} = 0.5$ | 1.7 (0.2) | 1.6 (0.4) |
| $b^{(2)} = 1$ | 1.9 (0.2) | 1.6 (0.3) |
| $b^{(2)} = 1.5$ | 1.9 (0.1) | 1.6 (0.2) |
| single trait expectation [Polechová and Barton, 2015] | | $b^{(1,m)} \approx 1.65$ |

Table S1: Maximal steepness at the margins of the realised range, indicating the critical steepness beyond which adaptation fails. We show the sum of both gradients $b^{(1,m)} + b^{(2)}$ at the realised margins (*i.e.* deme $m$, defined as the deme where $v^{(1,m)} + v^{(2,m)}$ is maximal; covariances were ignored here, as their relative contribution to the maximal variance was negligible) for the two-trait model and the composite single-trait model. Values were calculated by first determining where the maximum of the sum of adaptive variances is attained in each replicate, and measuring steepness of the composite gradient in that deme (on both sides), then averaging over the two to obtain an estimate for each replicate. The standard deviation is the uncorrected standard deviation (square root of the sample variance) over 50 replicates, each with two margins. The last row is the maximal steepness at which the population is expected to be able to adapt in the single steepening gradient model [Polechová and Barton, 2015].

| steepness $b^{(2)}$ | range extent [mean (SD)] | |
|---|---|---|
| | two traits (optima $\theta^{(1,i)}$ and $\theta^{(2,i)}$) | single trait (optimum $\theta^{(1,i)} + \theta^{(1,i)}$) |
| $b^{(2)} = 0$ | 84 (3) | 84 (3) |
| $b^{(2)} = 0.5$ | 72 (2) | 70 (4) |
| $b^{(2)} = 1$ | 62 (2) | 52 (5) |
| $b^{(2)} = 1.5$ | 42 (2) | 20⋆ (6) |

⋆ as shown in fig. 5, total genetic variance in the case of $b^{(2)} = 1.5$ is approximately constant over the realized range, which makes the estimate of the location of the deme where genetic variance is maximal less precise.

Table S2: Mean realised range extent, defined as the distance between the left margin and right margin, (defined as the deme $m_r$ or $m_l$ where $v^{(1,m_{l/r})} + v^{(2,m_{l/m})}$ is maximal; in the equation, $m_{l/r}$ is either the left or right margin), for the two-trait model and the composite single-trait model. Covariances were ignored here, as their relative contribution to the maximal variance was negligible. Means were calculated over 50 replicates. The standard deviation is the uncorrected standard deviation (square root of the sample variance). The last row is the maximal steepness beyond which the population is expected to fail to adapt in the single steepening gradient model [Polechová and Barton, 2015].

##### Average growth rate in the single-trait model

We repeat the calculations performed in Appendix B for the growth rate in the two-trait model, but here assuming that all loci contribute to a single adaptive trait. To distinguish the notations in the composite single-trait model from those used in the two-trait model, we use $r_{l,\tau}^{(c,i)}$ to denote the growth rate at generation $\tau$, for individual $l$ in deme $i$, for the composite single-trait model. We find that the population average growth rate ($\bar{r}_\tau^{(c,i)}$) can be expressed in terms of conveniently chosen phenotypic contributions $z^{(1,i)}$ and $z^{(2,i)}$ underlain by a separate set of loci, as in the two-trait model, as

$$\bar{r}_\tau^{(c,i)} = r_{\max}\left(1 - \frac{N_\tau^{(i)}}{K}\right) - \frac{\left(\bar{z}_\tau^{(1,i)} - \theta^{(1,i)}\right)^2}{2V_s} - \frac{\left(\bar{z}_\tau^{(2,i)} - \theta^{(2,i)}\right)^2}{2V_s} - \frac{\mathrm{Var}(z_\tau^{(1,i)})}{2V_s} - \frac{\mathrm{Var}(z_\tau^{(2,i)})}{2V_s} - \Delta\bar{r}_\tau^{(i)}\ , \quad \text{(S6)}$$

where

$$\Delta\bar{r}_\tau^{(i)} = \frac{\mathrm{E}\big((z_{\tau,l}^{(1,i)} - \theta^{(1,i)})(z_{\tau,l}^{(2,i)} - \theta^{(2,i)})\big)}{V_s}\ , \quad \text{(S7)}$$

is the difference between the growth rate for the two-trait model and the single-trait model. Thus, the population average growth rate in the two-trait model is by $\Delta\bar{r}_\tau^{(i)}$ larger than in the single-trait model. Note that, in the single-trait model, $\Delta\bar{r}_\tau^{(i)}$ is the covariance between the phenotype components $z^{(1,i)}$ and $z^{(2,i)}$ underlain by separate sets of loci. In other words, $\Delta\bar{r}_\tau^{(i)}$ is generated by the LD between these two sets of loci.

#### Decrease in growth rate due to deviation from the optimum

Here we show that the effect on individuals' fitness, due to a given deviation from the optimum, differs in the single- and two-trait model. We start by considering an individual such that, in the two-trait model, its phenotypes $z^{(1,i)}_{\text{two}}$ and $z^{(2,i)}_{\text{two}}$ deviate from the optimal phenotypes $\theta^{(1,i)}_{\text{two}}$ and $\theta^{(2,i)}_{\text{two}}$ by $\delta z^{(1,i)} = z^{(1,i)}_{\text{two}} - \theta^{(1,i)}_{\text{two}}$ and $\delta z^{(2,i)} = z^{(2,i)}_{\text{two}} - \theta^{(2,i)}_{\text{two}}$ (subscript $l$ is omitted for clarity). In the composite single-trait model, the deviation of the composite phenotype $z^{(1,i)}_{\text{two}} + z^{(2,i)}_{\text{two}}$ of this individual from the composite optimal phenotype $\theta^{(1,i)}_{\text{two}} + \theta^{(2,i)}_{\text{two}}$ is equal to $\delta z^{(c,i)}_{\text{one}} = z^{(1,i)}_{\text{two}} + z^{(2,i)}_{\text{two}} - \theta^{(1,i)}_{\text{two}} - \theta^{(2,i)}_{\text{two}} = \delta z^{(1,i)} + \delta z^{(2,i)}$. We find that the fitness of this individual is smaller in the single-trait model than in the two-trait model. Indeed, in the single-trait model, the individual difference in growth rate $\mathcal{L}_{\text{one}}$ due to deviations in the single-trait model is

$$\mathcal{L}_{\text{one}} = \frac{1}{2V_s}\left[(\delta z^{(1,i)})^2 + (\delta z^{(2,i)})^2 + 2\delta z^{(1,i)}\delta z^{(2,i)}\right];. \tag{S8}$$

By contrast, the corresponding decrease in growth rate in the two-trait model, $\mathcal{L}_{\text{two}}$, is

$$\mathcal{L}_{\text{two}} = \frac{1}{2V_s}\left[(\delta z^{(1,i)})^2 + (\delta z^{(2,i)})^2\right] . \tag{S9}$$

The difference in individual fitness costs between the two-trait and the single-trait model, helps explain why at the margins larger total genetic variance is sustainable in the two-trait model with respect to the single-trait model. Because the cost of fitness due to deviations from the optimum is larger in the single-trait model, individuals with phenotype that is too far from the optimum are less likely to survive and/or reproduce. In figure S8 we show the phenotypic component of the fitness (calculated per individual as $2\exp\sum_j -(z_l^{(j,i)} - \theta^{(j,i)})^2/(2V_s)$ and then averaged over the population of each deme). These patterns are reflected in larger population sizes at the margins in the two-trait model (fig. 4).

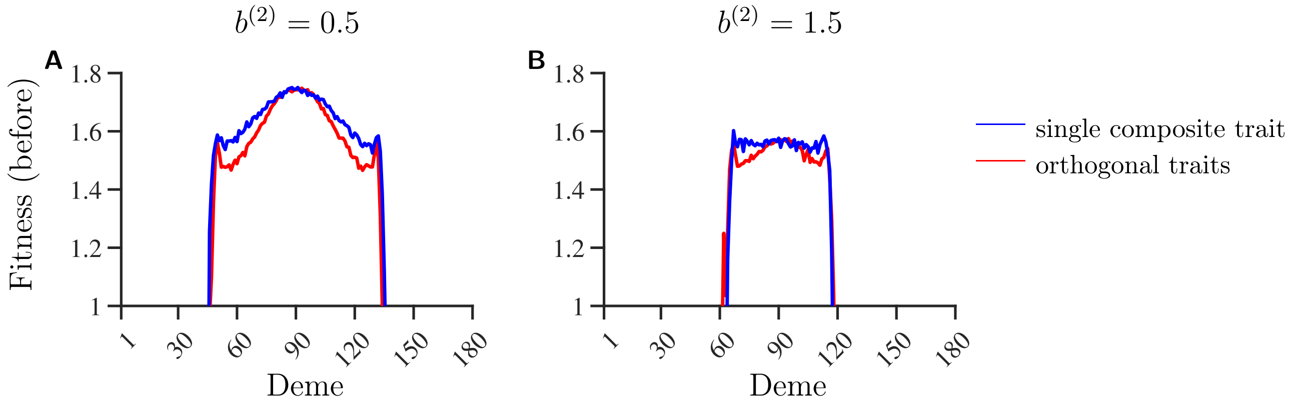


Figure S8: Comparison of the phenotype component of fitness between a single-trait model with single composite gradient $\theta^{(c,i)}$ (blue line), and two-traits model (red line). Panels (A), (B): phenotypic component of the fitness measured before migration and after selection, averaged over 50 independent realisations for different values of the steepness of the constant gradient, $b^{(2)} = 0.5$ (A) and $b^{(2)} = 1.5$ (B), as a function of deme position. Other parameters: $\alpha = 1/\sqrt{10}$, $V_s = 2$, $\sigma = 1$, $r_{\max} = 1$, $L = 653$, $K = 100$, and $\theta^{(1,i)}$, $\theta^{(2,i)}$ as given in Appendix A, (S1) and (S2) respectively.
